# Anoxygenic Photosynthesis and Dark Carbon Metabolism under micro-oxic conditions in the Purple Sulfur Bacterium “*Thiodictyon syntrophicum*” nov. strain Cad16^T^

**DOI:** 10.1101/420927

**Authors:** Samuel M Luedin, Nicola Storelli, Francesco Danza, Samuele Roman, Matthias Wittwer, Joël F Pothier, Mauro Tonolla

## Abstract

The microbial ecosystem of the meromictic Lake Cadagno (Ticino, Swiss Alps) has been studied intensively to understand metabolic functions driven by the highly abundant anoxygenic phototrophic sulfur bacteria of the families Chromatiaceae and Chlorobiaceae. It was found that the sequenced isolate “*Thiodictyon syntrophicum*” nov. sp. str. Cad16^T^, belonging to the Chromatiaceae, may fix 26% of all bulk inorganic carbon in the chemocline at day and night. With this study, we elucidated the mode of dark carbon fixation of str. Cad16^T^ with a combination of long-term monitoring of key physicochemical parameters with CTD, ^14^C-incorporation experiments and quantitative proteomics of *in situ* dialysis bag incubations of pure cultures. Regular vertical CTD profiling during the study period in summer 2017 revealed that the chemocline sank from 12 to 14 m which was accompanied by a bloom of cyanobacteria and the subsequent oxygenation of the deeper water column. Sampling was performed both day and night in September. While CO_2_ assimilation rates were higher during the light period, the relative change in the proteome (663 quantified proteins) was only 1% of all CDS encoded in str. Cad16^T^. Oxidative respiration was thereby upregulated at light, whereas stress-related mechanisms prevailed during the night. These results indicate that the low light availability due to high cell concentrations and the oxygenation of the chemocline induced a mixotrophic growth in str. Cad16^T^.

The complete proteome data have been deposited to the ProteomeXchange with identifier PXD010641.

## 2. INTRODUCTION

Inorganic carbon, nitrogen and sulfur is cycled in diverse microbial metabolism networks in paired redox reactions and organic compounds are thereby produced in the scale of 10^9^ t year^-1^ worldwide [1]. Light has been used as a source of energy in phototrophic anoxygenic bacteria since at least 3.85 Ga (billion years before the present) [2]. During carbon fixation, the transfer of electrons along a gradient of carrier molecules with sequentially lower potential allows the generation of energy bound to phosphate (ATP) and reductants [e.g., NAD(P)H and reduced ferredoxin]. The electrons required to replenish the oxidized electron acceptor pool can be derived from the oxidation of reduced sulfur species such as sulfide, sulfite and thiosulfate, or H_2_ and Fe (II) [3].

The anoxygenic photosynthetic purple sulfur bacteria (PSB) of the family Chromatiaceae are found widespread in aquatic, sulfidic oxygen minimum-zones where light is still available [4]. As an adaption to low light availability around 0.1–20 *µ*mol m^2^ s^-1^ and only limited wavelength (450–600 nm, and infrared above 750 nm), PSB contain pigments of the carotenoid and bacteriochlorophyll *a* and *b* (BChl) classes [5], as well as multiple copies of antenna peptides (LHC), to subtly modulate charge separation within the membrane bound type II reaction center (RC) [6, 7]. Carbon is typically fixed through the Calvin-Benson-Bassham (CBB) cycle [8]. In order to store both, reduction-equivalents and oxidized carbon, PSB intracellularly concentrate elemental sulfur-chains (S-S_n_^0^) in protein covered globules (SGBs) [9] and glycogen and polyhydroxybutyrate (PHB) [10], respectively. These assimilates may subsequently allow for chemotrophic growth in the dark [11, 12].

The possibility of chemolithoautotrophic growth of PSB under microaerophilic conditions in the dark has been proposed by van Niel [13] and has been described first for *Thiocapsa roseopersicina* BBS [14]. The PSB *T. roseopersicina* inhabiting shallow tidal flats is especially adapted to the daily changes of oxygen concentrations and competes with chemotrophic non purple sulfur bacteria (*Thiobacillus* spp.) and *Beggiatoaceae* spp. [15]. Chemoheterotrophic, chemoautotrophic and mixotrophic growth has since been shown for different PSB spp. [16–21]. Several strains of *Allochromatium vinosum* and *T. roseopersicina* have shown mixotrophic growth under a 5% oxygen atmosphere and acetate, and different reduced sulfur compounds [18].

The ecological significance and the impact on biogeochemistry of PSB were extensively studied in permanently stratified lakes [22–27]. In Lake Cadagno (Piora valley, Swiss Alps), underwater springs in gypsum rich dolomite provide a steady inflow of solute-rich water. In combination with solute-poor surface water, a stable and steep gradient in redox potential, salinity, sulfide and oxygen concentrations at around 12 m depth is formed [28]. Within this chemocline, a dense population of phototrophic sulfur oxidizing bacteria (PSOB) of the family Chromatiaceae and Chlorobiaceae (GSB; green sulfur bacteria) thrive to dense populations with up to 10^7^ cells ml^-1^ between June and October [29]. *In situ* chemocline incubation experiments with ^14^C-uptake in Lake Cadagno [30, 31] and other lakes [32], as well as with nanoSIMS ^13^C-stable-isotope labelling [33] revealed both light-driven and dark carbon fixation of PSB. Thereby it was found, that the population of PSB isolate “*Thiodictyon syntrophicum*” sp. nov. str. Cad16^T^ (str. Cad16^T^, thereafter) [34] assimilated 26% of the total carbon the chemocline during dark and light incubations [31]. Alternatively, the relative contribution of different PSB and GSB spp. to total carbon assimilation normalized to biomass during daytime was estimated with stable isotope analysis for Lake Cadagno. Thereby, str. Cad16^T^ only photosynthetically fixed 1.3 to 2 % of the carbon, as estimated from the daily bulk *δ*^13^C-mass balance [35].

Additional insight came from an *in vitro* quantitative proteomics study with str. Cad16^T^ growing anaerobically under light and dark with 1 mM H_2_S [36]. Photosynthesis-driven growth of str. Cad16^T^ resulted in the relative >1.5× expression of 22 proteins. Most notably, the poly(R)-hydroxyalkanoicacid synthase subunit PhaE and the phasin (PhaP) involved in the synthesis of PHB were found. In contrast, among the 17 proteins overexpressed under dark conditions, three enzymes of the dicarboxylate/4-hydroxybutyrate (DC/HB) cycle were detected, indicating dark carbon fixation through this typically *archeal* pathway.

The complete genome of str. Cad16^T^ gave evidence of the biological functions encoded [37]. Similar to *Allochromatium* spp. or *Lamprocystis* spp., str. Cad16^T^ expresses a type II (quinone type) RC, the membrane-bound protein cascade of cyclic electron transport to generate ATP and reverse electron transport to produce NAD(P)H, and also contains a *cbb*3 type cytochrome *c*. The later may enable microaerobic growth and Fe(III) oxidation of str. Cad16^T^ [38]. However, no genetic evidence for the possible syntrophic relationship within aggregates of *Desulfocapsa* sp., and also only incomplete Sox and no thiosulfate dehydrogenase Tsd proteins, responsible for SO_3_^2-^ oxidation, as previously described for str. Cad16^T^ by Peduzzi and colleagues [34], were found.

With this study, we aimed at elucidating the metabolic pathways of PSB str. Cad16^T^ in more detail, to better understand the key metabolic mechanisms involved. The main objectives of this study were: (i) to study differences between day and night in the metabolism of PSB str. Cad16^T^ *in situ* at the chemocline of Lake Cadagno and (ii) to monitor the environmental factors longitudinally that determine the metabolic activity of the chemocline community. We used a combination of CO_2_ assimilation analysis using ^14^C-scintillation and proteomics using label-free quantitation tandem mass spectrometry (LFQ-MS^2^) to quantify metabolic activity and the pathways involved in str. Cad16^T^. Relative light intensity and temperature at the depth of the chemocline were measured constantly and several CTD profiles were taken during the incubation. The carbon assimilation rates and protein profiles obtained, thereby revealed a mixotrophic metabolism of str. Cad16^T^ influenced by the unique microaerobic *in situ* conditions encountered in summer 2017.

## 3. MATERIALS AND METHODS

### 3.1. Study Site and Field Measurements

The *in situ* incubations were performed in Lake Cadagno between 13 July 2017 to 23 September 2017 with dialysis bags attached to a mooring (46°,33’,05,1” N/8°,42’,43,0” E, max. depth 18 m) (**suppl. Figure S 1b and c**). From 13 July to 13 September 2017 different physical and chemical parameters were measured alongside the incubations in order to monitor the *in situ* chemocline conditions and adjust the incubation depth, if necessary. A CTD (Conductivity, Temperature, Depth) 115 probe (Sea & Sun Technology GmbH, Germany) equipped with temperature, salinity, oxygen, redox potential, chlorophyll *a* (Chl *a*) and turbidity sensors was used to measure physicochemical profiles. Profiles from 13 July 2017 were taken as an estimate of chemocline depth (**suppl. Figure S2** HOBO UA-002-64 Pendant data loggers (Onset Computer Corporation, MA, USA) measured relative light (Lux; 180–1’200 nm) and temperature at 60 min intervals. Two sensors were placed near the surface (0.05 m depth) and other pairs were positioned 0.4 m apart at the upper and lower part of the rig, respectively (**suppl. Figure S 1c**). An empirical conversion factor of Lux = 0.018 PAR and 0.016 PAR (*µ*mol m^-2^ s^-1^) was used for the surface and below the water, respectively as in ref. [39]. The passive HOBO logger values were analyzed after retrieval at the end of the experiment. We additionally had access to CTD data from a parallel project form Dr. Oscar Sepúlveda Steiner and colleagues from EAWAG (Dübendorf, Switzerland) where two CTD profiles were taken daily.

### 3.2. Flow Cytometry for Cell Counting

Flow-cytometry based cell counting was performed as in Danza and colleagues [40]. In short, phototrophic bacteria were identified using 50 *µ*l sub-samples in triplicates with a BD Accuri C6 cytometer (Becton Dickinson, San José, CA, USA). A forward scatter threshold of FSC-H 10’000 was applied to exclude abiotic particles. A second red fluorescent (FL3-A) threshold above 1’100 was applied to select for cells emitting autofluorescence due to Chl and BChl. The flow rate was set to 66 *μ*l min^-1^.

### 3.3. Estimates of Biovolume and Biomass of Bacterial Cells

The biovolume was calculated for strain Cad16^T^ assuming a median diameter of 2 *µ*m (range; 1.4–2.4 *µ*m) for a spherical cell. Biomass was estimated using the conversion factor determined for Lake Cadagno PSB strains 4.5 fmol C m^-3^ [33].

### 3.4. Estimates of Carbon Uptake Rates

We took the estimate from Camacho *et al.* [30] that half of the carbon is fixed through oxygenic and anoxygenic photosynthesis, respectively. Musat and colleagues estimated anoxygenic carbon assimilation for PSB *C. okenii* and GSB *C. clathratiforme* to be 70% and 15% of the total daily CO_2_ assimilation, respectively [33]. We therefore calculated the uptake rates for the three populations as follows (Eq. 1) where *A*_day_ is the total uptake rate per cell of the phototrophic community and *F* the fraction as estimated in [33]:

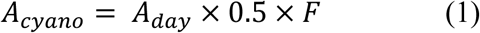

### 3.5. Bacterial Pure Cultures and Media

Str. Cad16^T^ was isolated in 2003 [41] and was subsequently grown in pure culture on Autotrophic Pfennig’s medium II [42] at the Laboratory of Applied Microbiology in Bellinzona, Switzerland (LMA) (**Figure S 1**a). The medium contained 0.25 g of KH_2_PO_4_ l^-1^, 0.34 g of NH_4_Cl l^-1^, 0.5 g of MgSO_4_·7H_2_O l^-1^, 0.25 g of CaCl_2_·H_2_O l^-1^, 0.34 g of KCl L^-1^, 1.5 g of NaHCO_3_ l^-1^, 0.02 mg of vitamin B12 l^-1^ and 0.5 ml of trace element solution SL12 l^-1^. The medium was autoclaved under a 80% N_2_ / 20% CO_2_ atmosphere [43] and 1.1 mM Na_2_S·9H_2_O was added aseptically. The pH was adjusted to 7.0. Cultures were grown in 500 ml glass bottles at ambient temperature and under a 12/12 h light/dark-regime with a 60 W incandescent lamp (6 *µ* mol quanta m^-1^ s^-1^). Cells were grown up to a concentration of around 3 × 10^6^ cells ml^-1^. Cell concentrations repeatedly were measured by flow cytometry.

Cellulose dialysis bags with a 14 kDa cutoff (D9777-100FT, Sigma-Aldrich, Buchs, CH) were rinsed for 1.5 h in Na_2_CO_3_ (40 g l^-1^) and 0.01 M EDTA at 60 °C wile stirring. The bags were cleaned with ddH_2_O, cut into 0.6 m long pieces, closed by a knot on one end and autoclaved for 20 min at 121 °C. On site, about 80 ml of bacteria culture were filled randomly in each bag, which was closed, attached to a rig and installed in the chemocline within 30 min (**suppl. Figure S 1b)**. In total 18 dialysis bags where placed at 12 m depth from 13 July to 23 August 2017 and then lowered to 14 m for the remaining campaign.

### 3.6. ^14^C-Incubations

The scintillation experiment was performed as in Storelli *et al.* [31]. In short, subsamples from three dialysis bags were pooled together randomly, 7 ml NaH^14^CO_3_ (NaH^14^CO_3_; 1.0 mCi; 8.40 mCi mmol^-1^, 20 *µ*Ci ml^-1^; Cat. No. NEC-086S Perkin-Elmer, Zurich, Switzerland) were added and incubated in cleaned and autoclaved 50 ml translucent Duran glass bottles (SCHOTT AG, Mainz, Germany). Six replicates of str. Cad16^T^ cultures were incubated for 4 hours during day (1:00–4:00 pm) and night (9:00 pm–12:00 am), respectively. Chemocline background fixation rates were determined in 50 ml chemocline samples. Filtered chemocline lake water (0.45 *µ*m) was used as negative control. Upon retrieval, the amount of *β*-activity (^14^C) assimilated by microbes during the incubation time was measured in the laboratory following standard method that included acidification and bubbling of the samples [44].

The inorganic dissolved carbon concentration was determined with the CaCO_3_ *Merck Spectroquant* kit No. 1.01758.0001 and the *Merck spectroquant Pharo 100* photospectrometer (Merck & Cie, Schaffhausen, Switzerland). Samples were taken at 14 m depth, filtered with 0.45 *µ*m filters, pH was tested with indicator paper (MColorpHast, Merck KGaA, Germany) to lie within 6.8–7.0 and triplicate samples were measured.

Scintillation was done on a *Guardian 1414* liquid scintillation counter (Perkin Elmer Wallac, MA, USA) running with the WinSpectral v.1.40 software. Raw data was statistically analyzed using *t*-tests in Excel (Microsoft Office 2010, v-.14.0.7168.5000).

### 3.7. Protein Extraction and Digest

Subsamples were transferred to 50 ml tubes upon retrieval and stored at 4 °C in the dark. They were then brought to the lab within 30 min and centrifuged 10’000 *g* at 4 °C for 10 min. The supernatant was discarded and the pellets were re-suspended in 1× PBS pH 7.0 and 1% EDTA-free Protease Inhibitor Cocktail (*v/v*; Thermo Fisher Scientific, Rheinach, Switzerland) and frozen at −20 °C until further processing.

The cells were thawed, lysed in 5% SDC (w/w) in 100 mM ammonium-bicarbonate buffer containing 1% EDTA-free Protease Inhibitor Cocktail (*v/v*; Thermo Fisher Scientific, Rheinach, Switzerland) and sonicated for 15 min at 200 W at 10 °C with a Bioruptor ultrasonicator (Diagenode SA, Belgium). Samples were then shipped to the Functional Genomic Center Zurich (FGCZ) on dry ice for further processing. Protein concentration was estimated using the Qubit Protein Assay Kit (Thermo Fisher Scientific, Rheinach, Switzerland). The samples were then prepared using a commercial iST Kit (PreOmics, Germany [45]) with an updated version of the protocol. Briefly, 50 *µ*g of protein were solubilized in ‘lyse’ buffer, boiled at 95 °C for 10 min and processed with High Intensity Focused Ultrasound (HIFU) for 30 s setting the ultrasonic amplitude to 85%. Then the samples were transferred to a cartridge and digested by adding 50 *µ*l of the ‘Digest’ solution. After incubation (60 min, 37 °C) the digestion was stopped with 100 *µ*l of Stop solution. The solutions in the cartridge were removed by centrifugation at 3’800 *g*, while the peptides were retained by the iST-filter. Finally the peptides were washed, eluted, dried and re-solubilized in ‘LC-Load’ buffer for Tandem Mass spectrometry (MS^2^)-analysis.

### 3.8. Liquid Chromatography and MS^2^-Analysis

MS^2^ analysis was performed on a *QExactive* mass spectrometer coupled to a nano *EasyLC 1000* HPLC (Thermo Fisher Scientific, Rheinach, Switzerland). Initial solvent composition was 0.1% formic acid for channel A and 0.1% formic acid, 99.9% acetonitrile for channel B, respectively. For each sample 4 *μ*L of peptides were loaded on a commercial Acclaim *PepMap* Trap Column (75 *µ*m × 20 mm; Thermo Fisher Scientific, Rheinach, Switzerland) followed by a *PepMap RSLC* C18 Snail Column (75 *µ*m × 500 mm; Thermo Fisher Scientific, Rheinach, Switzerland). The peptides were eluted at a flow rate of 300 nl min^-1^ by a gradient from 5 to 22% B in 79 min, 32% B in 11 min and 95% B in 10 min. Samples were acquired in a randomized order. The mass spectrometer was operated in data-dependent mode (DDA), acquiring a full-scan MS spectra (300-1’700 *m/z*) at a resolution of 70’000 at 200 *m/z* after accumulation to a target value of 3’000’000, followed by HCD (higher-energy collision dissociation) fragmentation on the twelve most intense signals per cycle. HCD spectra were acquired at a resolution of 35’000, using a normalized collision energy of 25 a. u. and a maximum injection time of 120 ms. The automatic gain control (AGC) was set to 50’000 ions. Charge state screening was enabled and singly and unassigned charge states were rejected. Only precursors with intensity above 8’300 were selected for MS^2^ (2% underfill ratio). Precursor masses previously selected for MS^2^ measurement were excluded from further selection for 30 s, and the exclusion window was set at 10 ppm. The samples were acquired using internal lock mass calibration on *m/z* 371.101 and 445.120.

### 3.9. Protein Identification and Label Free Protein Quantification

The acquired raw MS^2^ data were processed by MaxQuant v.1.4.1.2, followed by protein identification using the integrated Andromeda search engine. Each file was kept separate in the experimental design to obtain individual quantitative values. Spectra were searched against a forward str. Cad16^T^ database (6’237 coding genes), concatenated to a reversed decoyed *fasta* database and common protein contaminants (NCBI Assembly No. ASM281377v1; release date: 2017/12/07). Carbamidomethylation of cysteine was set as fixed modification, while methionine oxidation and N-terminal protein acetylation were set as variable. Enzyme specificity was set to trypsin/P allowing a minimal peptide length of 7 amino acids and a maximum of two missed-cleavages. Precursor and fragment tolerance was set to 10 ppm and 0.05 Da for the initial search, respectively. The maximum false discovery rate (FDR) was set to 0.01 for peptides and 0.05 for proteins. Label free quantification was enabled and a 2-min window for match between runs was applied. The re-quantify option was selected. For protein abundance, the intensity (Intensity) as expressed in the protein groups file was used, corresponding to the sum of the precursor intensities of all identified peptides for the respective protein group. Only quantifiable proteins (defined as protein groups showing two or more razor peptides) were considered for subsequent analyses. Protein expression data were transformed (hyperbolic arcsine transformation) and missing values (zeros) were imputed using the *missForest R*-package v.1.4 [46] The protein intensities were normalized by scaling the median protein intensity in each sample to the same values.

Scaffold v.4.8.4 (Proteome Software Inc., Portland, OR) was used to validate MS^2^ based peptide and protein identifications. Peptide identifications were accepted if they could be established at greater than 42.0% probability to achieve an FDR less than 0.1% by the Peptide Prophet algorithm with *Scaffold* [47] delta-mass correction. Protein identifications were accepted if they could be established at greater than 54.0% probability to achieve an FDR less than 1.0% and contained at least two identified peptides. Protein probabilities were assigned by the *Prophet* algorithm [48]. Proteins that contained similar peptides and could not be differentiated based on MS^2^ analysis alone were grouped to satisfy the principles of parsimony. Proteins sharing significant peptide evidence were grouped into clusters. For the two-group analysis the statistical testing was performed using (paired) *t*-test on transformed protein intensities (hyperbolic arcsine transformation). Proteins were called significantly differentially expressed if linear fold-change exceeded 2-fold and the *q*-value from the *t*-test was below 0.01.

As an alternative method to find differentially expressed proteins, we used the correlation adjusted *t*-Score algorithm provided by the *R*-package sda v.1.3.7 [49] to further analyze the dataset of 1’333 proteins identified with MaxQuant.

### 3.10. Protein Functional Annotation

BlastKOALA v.2.1 [50] and eggNOG v.4.5.1 [51] were used to classify the proteins into functional categories. The complete KEGG-dataset for str. Cad16^T^ can be found under Ref. [52].

### 3.11. Genomic Data Availability

The complete genome of str. Cad16^T^ [52] is available under the GenBank assembly GCA_002813775.1.

### 3.12. Proteomic Data Availability

The complete proteomic data of str. Cad16^T^ have been deposited to the ProteomeXchange Consortium [53] via the PRIDE partner repository [54] under the accession PXD010641 and project DOI 10.6019/PXD010641

## 4. RESULTS

### 4.1. Physicochemical Parameters from July to August 2017

Fluctuations in light intensity were positively correlated with the predicted surface radiation and negatively associated with cloud cover, respectively (**suppl. Figure S3 a**). Daily relative average surface light intensity at 12:00 pm was 1’448.9 *µ*mol quanta m^-2^ s^-1^ (4.3–3’769.6) (**suppl. Figure S3 b**). The surface temperature was stable at an average of 15 °C. High temperatures up to 40 °C were measured, due the solar heating of the logger casing (**suppl. Figure S3 b**). The accumulated sunlight were above average for the weeks observed from (**suppl. Figure S4**). From 13 July to 23 August 2017 at 12 m depth, an average of 3.7 *µ*mol quanta m^-2^ s^-1^ (0.2–26.2) was measured. At 12.4 m depth, a mean of 1.3 *µ*mol quanta m^-2^ s^-1^ (0.04–7.8) was registered (**suppl. Figure S3 c and d**). Within this period, changes of turbidity, oxygen and Chl-*a* profiles in the daily CTD profiles indicated that the chemocline had been sinking from 12 to 13–14.5 m depth (**suppl. Figure S5**). To ensure chemocline conditions for the incubations, we adjusted the depth of the rig from 12 to 14 m depth. This resulted in an unexpected reduced relative light intensity at the position of the rig at 14 m for the days 23 August to 13 September 2017. Only an average of 0.4 *µ*mol quanta m^-2^ s^-1^ (0.2–4.0) was measured at 14 m depth from 24 August to 13 September 2017, and no light was measured at 14.4 m depth after 23 August 2017. Temperature was stable around 5 °C (4.6–6.1) at both incubation depths with a positive trend over the months (**suppl. Figure S3 c and d**).

### 4.2. Chemical and Physical Analysis of Lake Cadagno at Sampling Date

Physicochemical measurements and endpoint-sampling was done on 12 and 13 September 2017. The weather was cloudless with weak wind. Two CTD profiles at 1:30 pm and 9:00 pm showed a comparable situation for temperature, dissolved oxygen and conductivity (**Figure 1**). Water temperature was stable at 12 °C from the surface down to the thermocline at 9.5 m, whereas it dropped to 5 °C at 16 m depth at both time points. Dissolved oxygen (DO) was measured at 8.5 mg l^-1^ (265.6 *µ*M) throughout the mixolimnion. From 7 m to 9.5 m depth, DO-concentrations steadily declined to 6 mg l^-1^, and then more rapidly to 2 mg l^-1^ at 10.5 m depth, at both time points measured. At 14 m depth, 0.4 mg l^-1^ (12.5 *µ*M) and 0.3 mg l^-1^ (9.4 *µ*M) were measured during day and night, respectively. Conductivity increased along the profile from 0.13 in the mixolimnion to 0.22 mS cm^-1^ hypolimnion, both at day and night. In contrast, a pronounced turbidity peak (30 FTU) was observed at a depth of 13 m at 1:30 pm whereas a broader distribution of the FTU values (6–16 FTU) from 13 to 14 m depth was observed at 9:00 pm. Water samples taken at 1:30 pm from 14 m depth showed a milky and pink coloration, characteristic for a concentrated PSB community. The total inorganic dissolved carbon concentrations measured at 14 m depth were 1.26 mM at 2:00 pm and 1.46 mM 9:00 pm.

**Figure 1.**
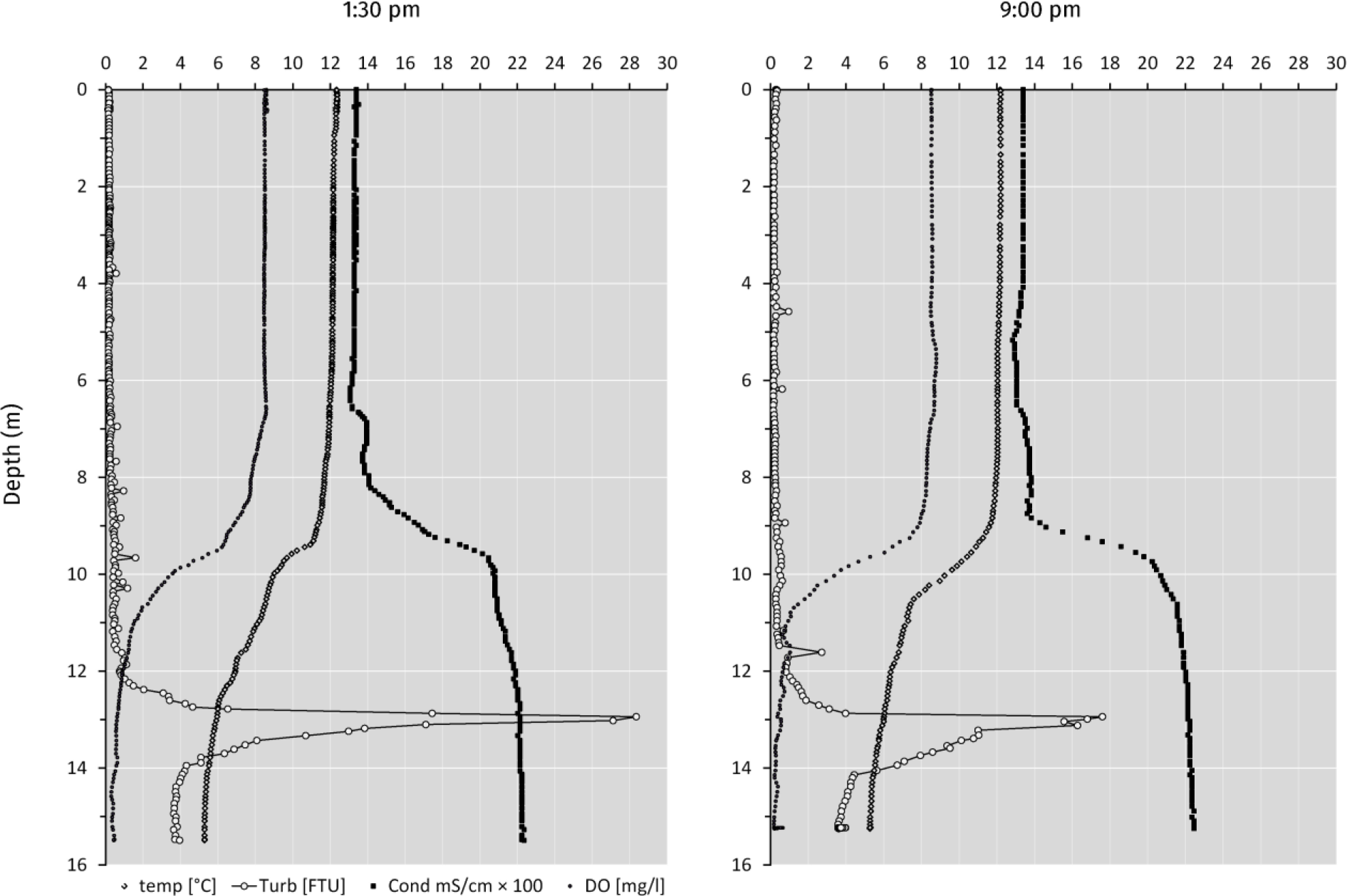
Conductivity, temperature, depth (CTD) profiles from Lake Cadagno on September 12 (1:30 pm) and (9:00 pm) 2017. Measurements were taken from the platform. Probe was equilibrated for 5 min at 0.5 m depth before measuring. Temperature (◊), FTU: Formazin Turbidity Unit (○), Cond: Conductivity (◼), DO: Dissolved Oxygen (.).

### 4.3. Microbial Counts and Evaluation

Initial str. Cad16^T^ cell concentrations were on average 3.1 × 10^6^ cells ml^-1^ in July, as measured by flow cytometry (FCM). The rigged cultures were checked on 23 August 2017 and all dialysis bags were found intact and cells were judged healthy due to the turbid-pinkish appearance. When retrieved for sampling on 12 September 2017, all dialysis bags were still intact, the population was uniformly distributed within the dialysis bags and the cells grew to a mean concentration of 9.3 × 10^6^ cells ml^-1^. No significant difference in cell concentration and internal complexity was measured between the two sampling groups (*P* = 0.74). In total, the str. Cad16^T^ cultures grew 3-fold from July to September.

The average cell concentration in the lake sample taken at 14 m was 4.23 × 10^6^ cells ml^-1^ at 1:30 pm, whereas it was 1.69 × 10^5^ cells ml^-1^ at 9:00 pm. However, the later value has to be questioned as the FCM count was below the values obtained for the 0.45 *µ*m filtered lake water of the ^14^C background control despite that the measured turbidity values were similar (4–5 FTU) at the two time points at 14 m sampling depth (**Figure 1**). FCM revealed that the phototrophic microbial community at 1:30 pm (4.23 × 10^6^ cells ml^-1^) consists mainly of *C. okenii*, *C. clathratiforme* and cyanobacteria spp. with 1.48 × 10^6^ cells ml^-1^, 7.45 × 10^5^ cells ml^-1^ and 1.49 × 10^6^ cells ml^-1^, representing 35, 35 and 17% of the total phototrophic population, respectively.

### 4.4. *In situ* Carbon Fixation Rates

Absolute carbon fixation rates at the chemocline were comparable between day and night when tested, with medians of 757 nM h^-1^ and 587 nM h^-1^, respectively (**Figure 2**). Cad16^T^ fixed carbon in both conditions of light or dark (**Figure 3**). For str. Cad16^T^ ^14^C-bicarbonate median uptake rates normalized per cell were significantly different between the conditions with 1’074 amol C cell^-1^ h^-1^ (range: 937–1’585) during the day, and 834 amol C cell^-1^ h^-1^ (range; 650–969) during the night (**Figure 3**). When the uptake rates were normalized to average biovolume (3.6 *µ*m^3^ cell^-1^), 316 amol C *µ*m^-3^ h^-1^ in light and 230 amol C *µ*m^-3^ h^-1^ in the dark were obtained, and a carbon-based doubling time range of 5 to 23 h was calculated for str. Cad16^T^. For the dominant phototrophic populations of the chemocline we estimated uptake rates in amol C cell^-1^ h^-1^ only at day because of uncertain FCM values at night. Thereby PSB *C. okenii*, GSB *Chlorobium* spp. and cyanobacteria assimilated in average 173.35, 73.70 and 245.68 amol cell^-1^ h^-1^, respectively.

**Figure 2.**
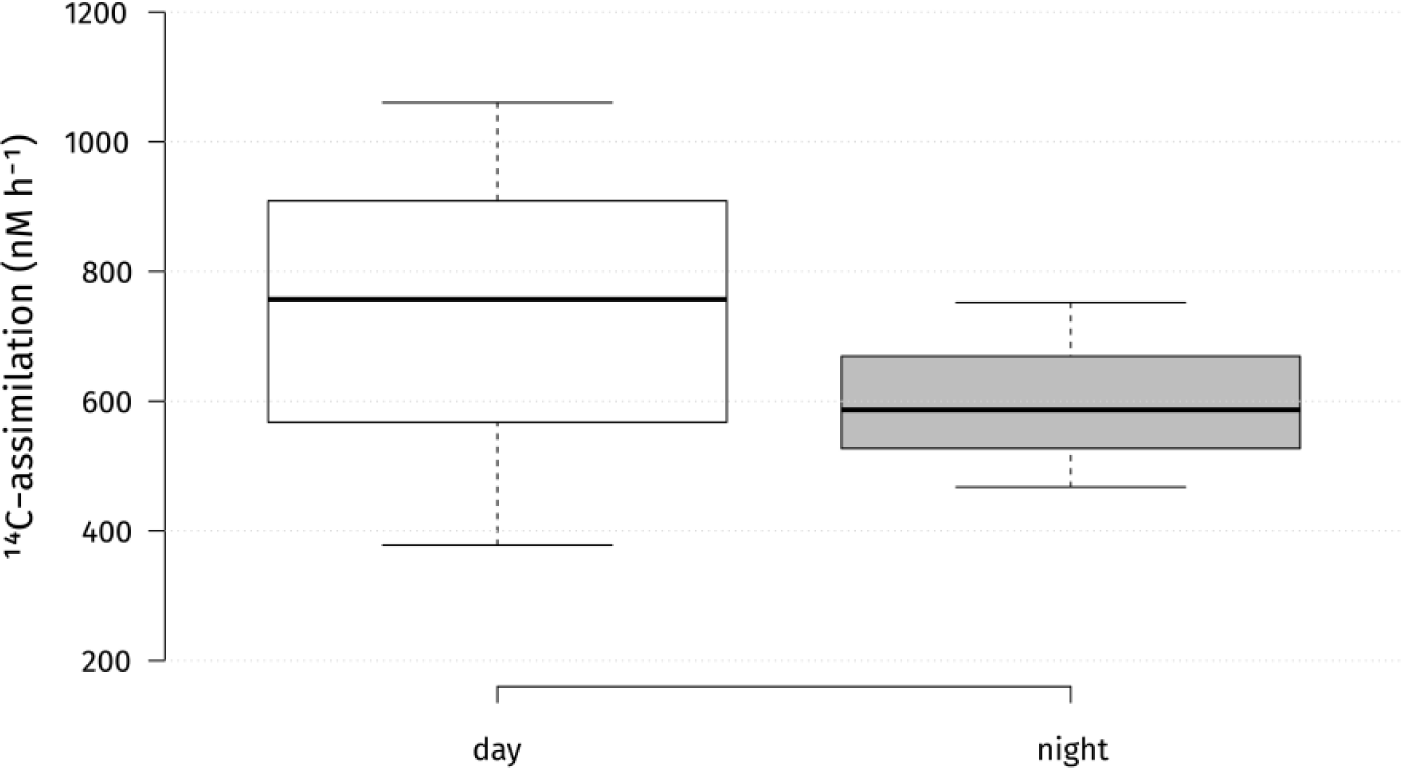
Absolute microbial ^14^C-uptake rates in the chemocline of Lake Cadagno during day and night. Center lines show the medians; box limits indicate the 25^th^ and 75^th^ percentiles as determined by *R* software; whiskers extend 1.5 times the interquartile range from the 25^th^ and 75^th^ percentiles, outliers are represented by dots. *n* = 3 sample points.

**Figure 3.**
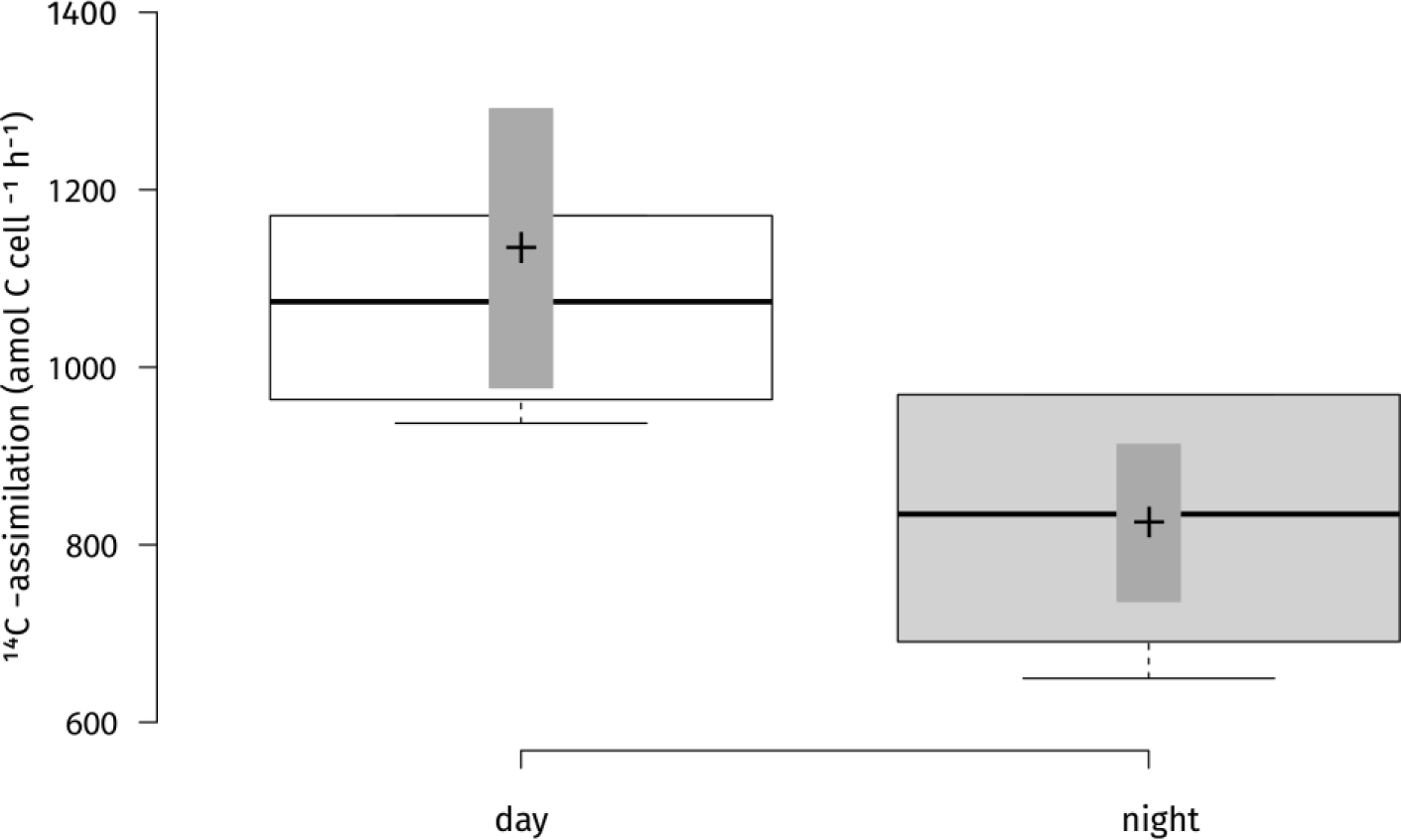
Carbon uptake rates per cell for strain Cad16^T^ cultures for two conditions (day /night) during 4 h incubation *in situ*. ^14^C-scintillation experiments were performed on six biological replicates. Two side *t*-test statistics was applied. The difference in uptake rates between the two conditions was statistically significant at *p* <0.05 (*p* = 0.02). Center lines show the medians; box limits indicate the 25^th^ and 75^th^ percentiles as determined by *R* software; whiskers extend 1.5 times the interquartile range from the 25^t^^h^ and 75^th^ percentiles, outliers are represented by dots; crosses represent sample means; bars indicate 83% confidence intervals of the means; data points are plotted as open circles. *n* = 6 sample points.

### 4.5. Total Proteins identified with LC-MS^2^-LFQ

A total of 11 samples, five samples for category ‘light’ and six samples for category ‘dark’, were processed and protein quantification was performed using the MaxQuant package. We used corrected *t*-test based statistics in order to identify and quantify proteins. The samples Cad16T_dia_7 (‘light’) and Cad16T_dia_12 (‘dark’) were identified as outliers in cluster analysis and were excluded from further data analysis. Therefore, the data analysis was made with four samples of the category ‘light’ and five samples of the category ‘dark’. Overall a total of 1’333 proteins (21% of the total coding CDS) with at least two peptides were identified.

### 4.6. Proteins Quantified with LC-MS^2^-LFQ

Peptide identifications were accepted if they could be established at >42.0% probability to achieve an FDR less than 0.1% with Scaffold delta-mass correction, resulting in 12’576 spectra included. Protein identifications were accepted if they could be established at >54.0% probability to achieve an FDR less than 1.0% and contained at least two identified peptides. The number of quantified proteins per condition was similar, with an average of 374 for “day” and 354 for “night”, respectively. Between 102 and 663 proteins were quantified in each biological replicate (7.2–37% of all IDs). Consequently, 684 proteins were quantified over all samples. Thereof, 21 contaminants were excluded. The remaining 663 CDS were classified with blastKOALA and EggNOG, with 627 annotated CDS for (99%) COG and 460 annotated CDS (69.4%) for blastKOALA, respectively.

As expected, many of the proteins with unchanged abundance belonged to the functional categories energy conversion, genetic information processing, carbohydrate and amino metabolism and protein modification (**Table 1**). Among the most abundant proteins detected were F_0_F_1_ ATPase subunits (AUB84561.1, AUB81565.1, AUB81565.1 and AUB84563.1) and chaperons (GroEL; AUB81575.1, AUB80010.1, AUB84066.1, DnaK; AUB84026.1). Since cell growth depends on protein synthesis, we found 36 ribosomal subunits as well as elongation factor Tu (AUB80476.1) to be equally abundant.

**Table 1.** Functional categories of “*Thyodictyon syntrophicum*” str. Cad16^T^ proteins quantified using. EggNOG v.4.5.1 **[50]** and blastKOALA v.2.1 [50]was used to classify proteins according to COG and KEGG classification, respectively.

| COG Code | Description | No. Entries |
| --- | --- | --- |
| J | Translation, ribosomal structure and biogenesis | 64 |
| K | Transcription | 15 |
| L | Replication, recombination and repair | 7 |
| D | Cell cycle control, Cell division, chromosome partitioning | 5 |
| V | Defense mechanisms | 3 |
| T | Signal transduction mechanisms | 19 |
| M | Cell wall/membrane biogenesis | 28 |
| N | Cell motility | 1 |
| U | Intracellular trafficking and secretion | 10 |
| O | Posttranslational modification, protein turnover, chaperones | 50 |
| C | Energy production and conversion | 93 |
| G | Carbohydrate transport and metabolism | 35 |
| E | Amino acid transport and metabolism | 52 |
| F | Nucleotide transport and metabolism | 12 |
| H | Coenzyme transport and metabolism | 27 |
| I | Lipid transport and metabolism | 18 |
| P | Inorganic ion transport and metabolism | 37 |
| Q | Secondary metabolites biosynthesis, transport and catabolism | 6 |
| **KEGG Functional Category** |  |  |
|  | Genetic information processing | 78 |
|  | Carbohydrate metabolism | 78 |
|  | Energy metabolism | 41 |
|  | Metabolism of cofactors and vitamins | 29 |
|  | Amino acid metabolism | 28 |
|  | Environmental information processing | 27 |
|  | Nucleotide metabolism | 17 |
|  | Metabolism of terpenoids and polyketides | 11 |
|  | Lipid metabolism | 9 |
|  | Metabolism of other amino acids | 5 |

The cells always contained the established components of the dissimilatory sulfate reduction pathway such as ATP-sulfurylase Sat (AUB82369.1), the adenylylsulfate reductase AprAB (AUB82371.1, AUB82370.1) and the sulfite reductase Dsr complex (AUB83448.1– AUB83455.1). A Sqr sulfide:quinone reductase and a glutathione amide reductase GarA homolog to *A. vinosum* putatively involved in intracellular sulfur shuttling [55] were also present. In PSB, sulfur oxidation provides the electrons for cyclic electron transport driven by light. Consequently, in str. Cad16^T^ PufMCL (AUB85378.1–AUB85380.1) and PuhA (AUB85431.1) subunits forming the RC II and six different PufAB antenna proteins (AUB85355.1–AUB85357.1, AUB85363.1, AUB85361.1, AUB85710.1) were expressed. Additionally we found enzymes for BChl synthesis, terpenoid backbone biosynthesis and carotenoid synthesis. Noteworthy, elements of the reduction pathways driven by photosynthesis are shared with oxidative phosphorylation in PSB. We found in total 23 protein subunits involved in substrate respiration including the NADH dehydrogenase subunits NuoCDEFG an HoxFU, the cytochrome reductase CytB and Cyt1, seven F-type ATPase subunits and two *cbb*3 cytochrome *c* oxidase subunits. PSB use the ATP and NAD(P)H derived from photosynthesis to fix CO_2_ through the CBB cycle. For str. Cad16^T^ a complete CBB cycle with the key enzymes CbbM/RbcL RuBisCO form II (AUB81831.1) and phosphoribulokinase PrkB (AUB79979.1) were present. The fixed carbon enters the central carbon metabolism as 3-phospho-D-glycerate. In both growth conditions, str. Cad16^T^ contains enzymes for glycolysis and gluconeogenesis, as well as pyruvate oxidation, the glyoxylate cycle and the citrate cycle (TCA cycle) in unvaried abundance. Additionally, the presence of malic enzyme (MaeB; AUB82893.1) may allow the entry of malate into the central carbon pathway via pyruvate, is shown for *A. vinosum*. In both conditions examined, the PHB synthase subunits PhaC and PhaE are expressed (AUB80707.1 and AUB84676.1). Also enzymes necessary for amino acid biosynthesis and Co-factor and vitamin synthesis were expressed under both groups analyzed.

In a second step, we analyzed the proteome data for significant changes between growth conditions. However, we found a large within-group variation and no proteins were significantly differently expressed between the two growth conditions using (paired) corrected *t*-test on transformed protein intensities (q.mod <0.01).

### 4.7. Proteins Differentially Expressed

The expression dataset was alternatively analyzed using correlation-adjusted *t*-scores (CAT scores) in order to additionally address the correlative structure of the dataset as only 4.5% of the proteins are differentially expressed. Thereby, 60 proteins were found differentially expressed (1% of all coding CDS) at a local false discovery rate of lfdr<0.05. (**Table 3**)

During the ‘day’ period 21 CDS were found relatively more abundant for str. Cad16^T^. Thereof, all CDS were annotated with eggNOG (**Table 2**). Growing in the light, str. Cad16^T^ over-expressed four proteins involved in oxidative phosphorylation that were subunits of NADH:quinone oxidoreductase, cytochrome *bc*1 complex respiratory unit and AtpC subunit of F-type ATPase. Two enzyme involved in the central carbon pathway were abundant, the glycogen synthase GlgA and the glucose-6-phosphate isomerase GPI involved in glycolysis. The enzyme acetolactate synthase I/III small subunit associated with thiamine synthesis and the A 7,8-dihydroneopterin aldolase FolB involved in tetrahydrofolate biosynthesis were additionally found. Also membrane transport systems were found, including the ABC-2 type transport system and Twin-arginine translocation (Tat). Additionally, chaperone-type proteins DnaJ and proteolytic ClpX were also over-expressed.

**Table 2.** List of “*Thiodictyon syntrophicum*” str. Cad16^T^ proteins identified as more abundant in the light period. Lfdr: local false discovery rate, cat: t-score for each group and feature the cat score of the centroid versus the pooled mean [79].

| Accession | Name | COG category | cat score | lfdr |
| --- | --- | --- | --- | --- |
| AUB81546.1 | cytochrome c1 | C | 5.44 | 9.53E-11 |
| AUB83791.1 | NADH-quinone oxidoreductase subunit I | C | 4.35 | 8.32E-05 |
| AUB84560.1 | F0F1 ATP synthase subunit epsilon | C | 3.51 | 2.26E-04 |
| AUB83997.1 | acetolactate synthase small subunit | E | 2.88 | 1.23E-02 |
| AUB81260.1 | glucose-6-phosphate isomerase | G | 3.63 | 2.26E-04 |
| AUB83261.1 | starch synthase | G | 4.48 | 2.53E-06 |
| AUB79823.1 | thiazole synthase | H | 2.74 | 2.93E-02 |
| AUB81699.1 | dihydroneopterin aldolase | H | 3.41 | 1.29E-03 |
| AUB83078.1 | DNA topoisomerase I | L | 2.98 | 3.81E-03 |
| AUB84659.1 | outer membrane lipoprotein carrier protein LolA | M | 5.43 | 2.03E-10 |
| AUB82245.1 | HflK protein | O | 6.34 | 1.24E-13 |
| AUB82609.1 | ATP-dependent protease ATP-binding subunit ClpX | O | 3.38 | 1.29E-03 |
| AUB84025.1 | molecular chaperone DnaJ | O | 6.17 | 1.24E-13 |
| AUB79510.1 | hypothetical protein | S | 2.84 | 1.23E-02 |
| AUB81403.1 | hypothetical protein | S | 3.53 | 2.26E-04 |
| AUB82197.1 | hypothetical protein | S | 3.54 | 2.26E-04 |
| AUB82687.1 | hypothetical protein | S | 3.96 | 1.21E-04 |
| AUB83592.1 | histidine kinase | S | 3.51 | 1.29E-03 |
| AUB84052.1 | host attachment protein | S | 3.51 | 2.26E-04 |
| AUB84690.1 | hypothetical protein | U | 3.02 | 3.81E-03 |
| AUB80231.1 | multidrug ABC transporter ATP-binding protein | V | 3.83 | 1.21E-04 |

**Table 3.**
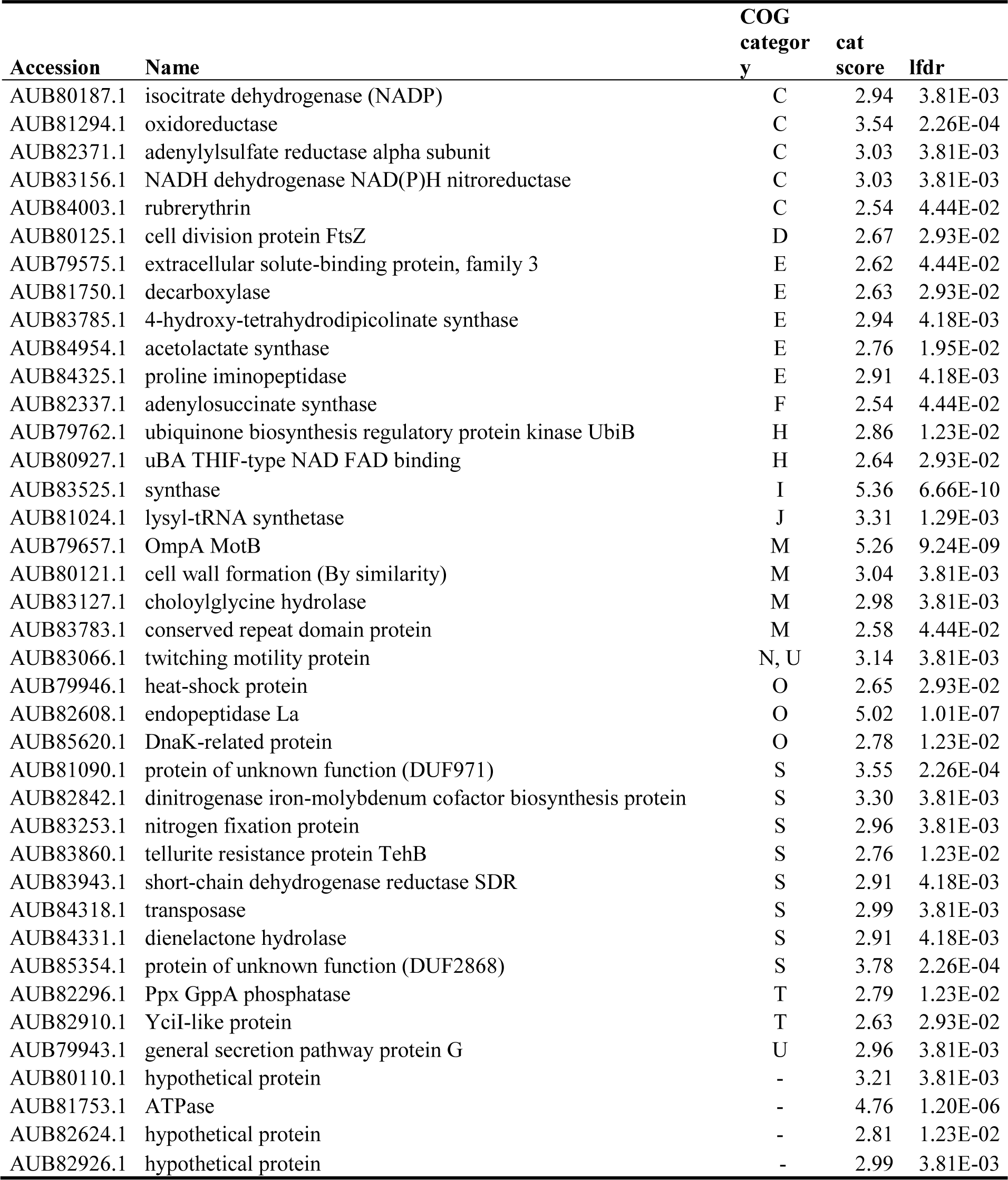
List of “*Thyodictyon syntrophicum*” str. Cad16^T^ proteins identified as more abundant in the dark period. Lfdr: local false discovery rate, cat: *t*-score for each group and feature the cat score of the centroid versus the pooled mean [79].

Under the condition ‘night’ 39 proteins were shown to be more abundant and thereof 35 entries (89%) were COG annotated (**Table 3**). Analysis of this data suggested that the NAD(P)H isocitrate dehydrogenase Idh1 responsible first carbon oxidation from oxaloacetate to 2-oxoglutarate in the TCA cycle as well as a FabH 3-oxoacyl-[acyl-carrier-protein] synthase III involved in fatty acid biosynthesis initiation and elongation were abundant. The adenylylsulfate reductase subunit alpha was found, and AprA is responsible for sulfite oxidation to 5’-adenylyl sulfate [56]. Cad16^T^ further expressed proteins associated to cell division (FtsZ), cell wall formation CpxP, Lysine and branched amino acid synthesis and nucleotide metabolism (Ppx-GppA; exopolyphosphatase and RutE; 3-hydroxypropanoate dehydrogenase). Elements of two secretion pathways were identified, a putative polar amino acid transport system and type II general secretion system. Additional three proteins detected to be more abundant in the dark are involved in stress response. The BolA-family transcriptional regulators, is a general stress-responsive regulator. Rubrerythrin may provide oxidative stress protection via catalytic reduction of intracellular hydrogen peroxide and a ATP-dependent serine protease mediates the degradation of proteins and transitory regulatory proteins, and thereby ensures cell homeostasis

## 5. DISCUSSION

We compared light to dark carbon fixation metabolism of PSB str. Cad16^T^ through a combination longitudinal monitoring of physicochemical condition of the Lake Cadagno chemocline, scintillation and quantitative proteomics of cultures incubated *in situ*.

### 5.1. Monitoring Data

During the incubation experiment from July to September 2017 the chemocline was actively monitored with CTD and light and temperature profiles were measured passively. This detailed record allowed us to understand the prevailing physicochemical conditions experienced by the str. Cad16^T^ cultures. Thereby, the average light availability measured was 10× lower than previously recorded at the chemocline, whereas the available DIC concentration was comparable to 2013 [31]. CTD measurements (this study) and FCM counts of the chemocline throughout the summer 2017 [57] revealed dense PSB and GSB populations (10^6^ cells ml^-1^) and an additional cyanobacterial bloom down to the monimolimnion from July 2017 on [40] that probably resulted in increased self-shading below a depth of 13 to 14 m [57]. As a consequence, the maximal peak of turbidity related to the PSOB sunk from around 12 to around 14 m depth. For this reason, we decided to hang the dialysis bags lower from 12 to 14 m at the 23 August 2017. The resulting low light conditions at 14 m depth possibly reduced net photosynthesis and subsequent carbon storage capacity and growth for str. Cad16^T^ as previously observed for PSB *Chromatium* spp. in the Lakes Cisó and Vilar [58, 59]. Whereas the monimolimnion has been typically described as anoxic [28], recurrent blooms of oxygenic plankton [30, 57] and optode-based measurements of dissolved oxygen has revealed micro-oxic conditions below 20 nM at the chemocline [60]. Our study helps to understand the consequences of these conditions for the PSOB.

### 5.2. Carbon Uptake Rates

Measured chemocline CO_2_-fixation rates at both conditions were within the ranges previously obtained for Lake Cadagno that were between 85 to 8’000 nM h^-1^ with light, and between 27 to 7’608 nM h^-1^ in dark conditions, respectively (**suppl.** Fehler! Verweisquelle konnte nicht gefunden werden.) [30, 33, 61, 31]. Comparable CO_2_-assimilation rates have been also measured in other stratified sulfureta as in Spanish karstic lakes with up to 3’000 nM h^-1^ both in light or dark conditions [62].Importantly, it was estimated that only half of the bulk carbon is fixed by anoxygenic photosynthesis in the Lake Cadagno chemocline [30]. Thereof, PSB *C. okenii* is responsible for 70% of the total anoxygenic phototrophic CO_2_ assimilation [33] and the concentration was 10^6^ cells ml^-1^—equal to 35% of the total phototrophic microbes— at 14 m depth in this study. Taken together, *C okenii* may have accounted for an average 173.35 amol C cell^-1^ h^-1^ in light conditions in our study, which is 60-fold lower that previously observed [33]. The average GSB assimilation rate with 73.7 amol C cell^-1^ h^-1^ was higher than observed (1–30), but within the same magnitude [33]. Oxygenic photosynthesis contributed 245.68 amol C cell^-1^ h^-1^ to the C-fixation that is 10–100 less than observed in the Baltic sea [63].

For str. Cad16^T^, the median uptake rate of 1’073.9 amol C cell^-1^ h^-1^ during the day was within the range measured for other PSB (100–30’000 amol C cell^-1^ h^-1^) [33, 31], however 10× lower compared to a previous *in situ* ^14^C-assimilation study with strain Cad16^T^ (around 12’000 amol C cell^-1^ h^-1^) [31]. In the former experiment, autotrophic CO_2_ assimilation rates of str. Cad16^T^ were additionally measured *in vitro*, with values between 8’541.7–18’541.7 amol C cell^-1^ h^-1^ with light, and around 2’916.7 amol C cell^-1^ h^-1^ in the dark, respectively. For C-fixation at night, highly variable C-assimilation rates in chemocline bulk samples were measured, ranging from 7–45% of the rates measured with light [24]. Schanz and colleagues thereby found a positive correlation between photosynthetically driven increase in overall biomass and dark fixation rates [24]. In our study, the median dark fixation rate of 834.4 amol C cell^-1^ h^-1^ was significantly lower than during the day, and again around 10× lower than measured by Storelli and colleagues [31]. This overall discrepancy may be explained by the 10-fold lower light availability that possibly reduced photosynthetic carbon assimilation and carbon storage (see above). Furthermore the sampling times in our study were adapted to the natural light-dark hours. This was in order to account for possible circadian effects, whereas in the former study incubations were performed in parallel at daytime, using clear and opaque sample bottles.

The differences between the study results may further be explained with the varied cell counting methodologies. Fluorescent *in situ*-hybridization (FISH) was used to count cells in the study of Storelli *et al.* [31]. Furthermore, we did not control for dead cells in FCM which might have led to a relative overestimation of the str. Cad16^T^ cell concentration. Moreover, str. Cad16^T^ forms aggregates, and some stuck to the dialysis bags visible as a pinkish residue, possibly reducing the number of planktonic cells for FCM counting.

When normalized to biomass, the carbon uptake values at light were higher as previously measured for PSB *C. okenii* and *Lamprocystis* sp. [33]. Uptake rates normalized to biovolume depend on the cell volume estimation (i.e. volume depends on r^3^) and the conversion factors chosen. We used values previously used in literature (see methods). For str. Cad16^T^, the estimated carbon-based doubling time was between 4 and 23 h, which is in contrast to previous estimates of 121 h *in vitro* [34], and a median of 333.6 h (25^th^ percentile; 196.8 h, 75^th^ percentile; 652.8 h) for the bulk biomass in the chemocline [24]. In contrast, the average doubling time obtained by FCM counting was 948.0 h (39.5 d) which is 40× longer than our ^14^C-uptake based calculations, and 8× longer than *in vitro* [34]. We did not control for the presence and metabolic activity of the syntrophic *Desulfocapsa* sp. nov. str. Cad626 [69] as it was observed that the str. Cad16^T^ *in vitro* cultures we used lost the initially co-occurring str. Cad626 after several years under laboratory growth conditions.

The significantly higher inorganic C-uptake rate during the day compared to night-time rates suggests active photosynthesis of str. Cad16^T^ at low light intensities, as it was described for other PSB [24, 64]. Noteworthy, the presence of up to 3 × 10^5^ ml^-1^ oxygenic phototrophic microbes down to 16 m depth [40] resulted in a partly oxygenated chemocline with around 0.6 mg O_2_ l^-1^ (19 *µ*M), as measured with CTD. Consequently, the chemocline waters retained some of the produced oxygen through the night (**Figure 1**). Taken together, str. Cad16^T^ may have used the O_2_ present as electron acceptor during mixotrophic growth under both conditions. As a consequence, some of the CO_2_ assimilated might have been constantly respired with thiosulfate as electron donor as found for *T. roseopersicina* str. M1 [15]. In accordance, str. Cad16^T^ grew in the dark with thiosulfate and 5% O_2_ *in vitro* [41]. Interestingly however, we found only SoxXY and SoxB and no TsdA thiosulfate-oxidizing enzyme homologues encoded in the str. Cad16^T^ genome [52]. As the complete Sox-complex is essential for the complete thiosulfate oxidation to sulfate in *A. vinosum* DSM 180^T^ [65], str. Cad16^T^ possibly uses an alternative mechanism. One alternative may be thiosulfate uptake through CysTWA (AUB80378.1, AUB80379.1 and AUB80380.1) and CysP (AUB80377.1) and oxidation to sulfite via the intermediate S-sulfocysteine by cysteine synthase B CysM (AUB82938.1) and possibly monothiol glutaredoxin of the Grx4 family (AUB83488.1) as suggested by Dahl [66]. However, with the applied methods we could not determine the relative contributions of photosynthetic or chemotrophic activity to the total increase in biomass. To conclude, ^14^C-fixation rates and FCM counting indicated an actively growing str. Cad16^T^ population during the incubation. The findings on carbon assimilation rates are further reflected in the proteome (discussed below).

### 5.3. Stable Fraction of the Strain Cad16^T^ Proteome

In order to understand differences in metabolism, the str. Cad16^T^ light/dark proteome has been studied extensively. Previously, Storelli and colleagues [36] performed a comparable proteomics study on light/dark metabolism, str. Cad16^T^ *in vitro* with 2D-difference gel electrophoresis (2D-DiGE), where they identified around 1’400 proteins, quantified 56 and, thereof, found 37 differentially expressed proteins. With six biological replicates instead of three, and the increased resolution and high-throughput capacity of LC-MS^2+^ compared to 2D-DiGE [67], we expected to substantially increase protein identification and quantitation in comparison. Overall, the number of identified proteins was comparable in both studies, with 1’400 in Ref [36] to 1’333 in our study. Noteworthy, LC-MS^2^ analysis returns sequence information on all proteins identified, whereas 2D-DiGE is limited to a chosen subset, based on the relative intensity and size of the protein spots on the gel. Therefore, we could quantify as much as 663 proteins that are 12× more than in the former study. Nevertheless, the number of unique proteins with different abundance between the two growth conditions was comparable, with 62 in our study versus 37 in the former, respectively. Interestingly, we retrieved only 28 of these proteins in the constantly expressed fraction of the proteome.

In addition to technical differences, the former study was performed *in vitro* under an anaerobic atmosphere, with higher light intensities (6 *µ*mol m^-2^ s^-1^) and at 20 °C, which may have an influence on metabolic rates and relative activity. In conclusion, the results of the present study can only be compared to some extend with previous findings.

The photosynthesis apparatus was abundant in both growth conditions, as several LHC proteins were detected. In accordance, also the enzymatic pathway for BChl*a* and carotenoid synthesis was expressed. In *T. roseopersicina* BChl*a* was absent at prolonged exposure to 60 *µ*M O_2_ in darkness [68]. In contrast, for str. Cad16^T^ we did not observe a loss of pigmentation. This might suggest that BChl*a* was not regulated on expression level and/or the micro-oxic conditions did not influence BChl*a* biosynthesis. The central role of dissimilatory sulfur oxidation during photosynthesis is well established for PSB [69] and proteins involved were found expressed in str. Cad16^T^. The proteins Sqr, Dsr and Sat were present and SGBs were also observed microscopically in both conditions. The CBB cycle is central in purple bacteria, not only for autotrophic carbon fixation, but also to regenerate the pool of reduced co-factors NAD[P]H_2_ [70]. Strain Cad16^T^ contains two forms of RuBisCo, RbcAB form I and RbcSL form II. In a previous study Storelli and colleagues [31] detected constitutive transcription of the *rbcL* gene under autotrophic condition *in vitro* under a 12/12 h dark/light regime, whereas the form I *rbcA* was induced by light. In contrast both *rbc* genes were transcribed equally under heterotrophic conditions with acetate, with and without light, respectively [31]. In our study we detected RbcL to be equally abundant in both conditions and no other RuBisCo subunits were found. Therefore, the sole presence of the dimeric form II RuBisCo may underline the importance of CO_2_ fixation mediated by the CBB cycle in maintaining the redox-balance under chemo or mixotrophic growth at low light and dark conditions, as described for purple bacteria [71].

In str. Cad16^T^, the CsrA (AUB84364.1) seems to be the main carbon storage regulator where it was detected under both conditions. Glycolysis under mixotrophic conditions might thereby be regulated through mRNA transcription and stability as in *A. vinosum* [56]. We additionally found enzymes involved in the central carbon pathways TCA, EMP and glyoxylate cycle in unvaried abundance. Noteworthy, isocitrate lyase was found expressed, that is involved in the glyoxylate cycle, that prevents loss of CO_2_ and ensures production of NAD[P]H_2_ otherwise occurring through the isocitrate dehydrogenase and 2-oxoglutarate dehydrogenase in the TCA [72]. Further, the malic enzyme was abundant, that generates oxaloacetate via malate through anaplerotic reactions without ATP [73]. The *ccb*3 cytochrome *c* oxidase found in both conditions is used in aerobic respiration and additionally used for FeS oxidation and it was speculated that str. Cad16^T^ is also involved in both aerobic [38] and anaerobic cryptic iron cycling, as found for *Thiodictyon* str. F4 [74]. Taken together, the metabolic fate of the carbon fixed is only partly elucidated by proteome analysis and metabolic carbon pathways may be active under the conditions examined.

### 5.4. Differentially Expressed Proteome

In light conditions, members of the oxidative respiration pathway were upregulated, indicating an active substrate respiration with light as in *T. roseopersicina* [20]. However, the cytochrome *bc* and the NADH-dehydrogenase complex and redox carrier molecules are also used in cyclic electron transport during photosynthesis in PSB. During light conditions, both chemotrophic and phototrophic metabolism compete for electrons in str. Cad16^T^. We also found evidence for glycolysis / gluconeogenesis since GPI was overexpressed. Interestingly, the glycogen synthase GlgA was additionally found abundant in the light. Glycogen synthesis and sulfur oxidation was found to be ineffectively regulated in *A. vinosum* [75] and we might speculate that in str. Cad16^T^ both processes are highly active even at slow growth, allowing for both, intracellular sulfur and glycogen accumulation. Altogether, these results indicate an active phototrophic and chemotrophic metabolism competing for electrons in the light.

In the dark, the micro-oxic condition may have led to the production of reactive oxygen species in str. Cad16^T^ that would explain the upregulated proteins involved in stress induced damage-control.

Interestingly, AprA was more abundant in the night period, indicating a relative higher activity of sulfite oxidation possibly coupled to chemotrophy. SGBs are consumed under dark autotrophic conditions in PSB *C. okenii* and *C. minus* observed *in vitro* [12], however the side scatter values measured did not vary between the two sample groups in this study when monitored with FCM. This may be due to glycogen and PHB storage inclusions, structures that add up additional structural complexity and depolymerization kinetics may be different to those of SGB. Supporting evidence is given by the unchanged presence of enzymes involved in of glycogen and PSB synthesis.

In summary, the 60 proteins found differentially expressed represent only about 1% of all protein coding CDS and about 5% of the identified proteins. Therefore, their impact on metabolic pathways is unclear and has to be further examined

## 6. CONCLUSION AND OUTLOOK

PSB str. Cad 16^T^ is metabolically flexible and growths phototrophically as well as chemotropically in the light as shown in this study. In dark conditions, low levels of oxygen may enable respiration of different small organic molecules. In order to understand the dark carbon metabolism, uptake experiments with labelled acetate and or pyruvate should be therefore included in the future. The long-term observation of the chemocline revealed the relative importance of oxygenic phototrophs on the oxygenation of the chemocline. However, it is not clear if O_2_ or reduced sulfur is used as terminal electron acceptor in str. Cad16^T^. Furthermore, the oxidation of Fe(II) to fix CO_2_ should be tested for str. Cad16^T^ *in vitro*, both under microaerobic and anaerobic conditions as found in Lake Cadagno in 2017. To better estimate the relative metabolic contributions of the different phototrophic energy metabolism at microaerobic conditions oxygenic photosynthesis-inhibitors [76] should be included in future experiments as a control. The contribution of phytoplankton to dark carbon uptake [77] has also not yet been elucidated in Lake Cadagno and therefore a sequential size dependent fractionation of sub-samples would be interesting. In order to further understand transcriptional control over the light to dark metabolism also mRNA sequencing experiments would be needed. To complete the understanding metabolomic studies would give insight into the relative amount of metabolic intermediates produced under different regimes. We further observed a large variability in the C-uptake rates between different studies that cannot be readily explained by differences in the chemocline community composition and/or technical variances. Therefore, NanoSIMS experiments [78] on Cad16^T^ may help to determine the within-population heterogeneity in inorganic C-assimilation under variable growth regimes.

## 8. AUTHOR CONTRIBUTIONS

SML, NS, FD, JFP, AB and MT conceived the study. SML, NS, FD and SR installed the mooring and performed field measurement and sampling. NS and FD prepared scintillation samples. FD did flow cytometry cell enumeration. SML extracted total protein and performed scintillation measurements. SML, AB, MW and JFP and MT analyzed physicochemical, proteomic and scintillation data. SML, NS; FD and MT prepared the manuscript. All authors contributed to writing and agreed on the manuscript before review.

## 9. ACKNOWLEDGEMENTS

We want to specially thank Michael Plüss (EAWAG) and Sébastien Lavanchy (EPFL) for the help in designing and installing the mooring. Furthermore, we are grateful to Damien Bouffard (EAWAG), Dr. Oscar Sepúlveda Steiner, Prof. Johny Wüest and Dr. Hannah Chmiel (EPFL) for sharing the physical datasets from their research. We are also thankful for the Angelo Carlino and Emilie Haizmann who made longitudinal measurements. For the LFQ-MS service and support in the analysis we want to thank Claudia Fortes and Jonas Grossmann from the FGCZ. We also thank the Alpine Biology Center Foundation (ABC) for their logistic support during fieldwork and Dr.Andreas Bruder for discussions and inputs during several stages of the study.

## APPENDIX

### Supplementary Tables

**Table S1.**
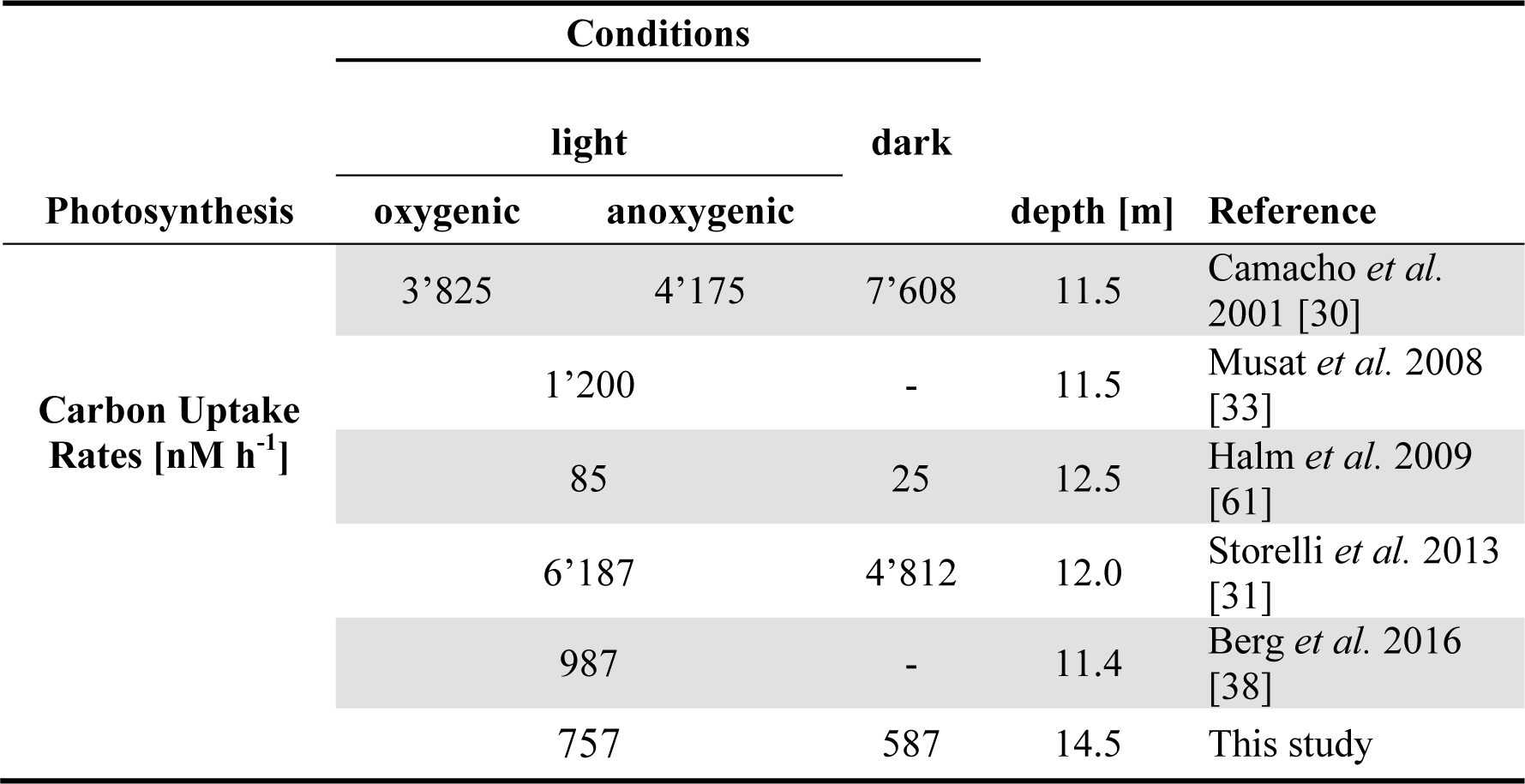
Absolute carbon uptake rates of the microbial community in the Lake Cadagno chemocline.

### Supplementary Figures

**Figure S1.**
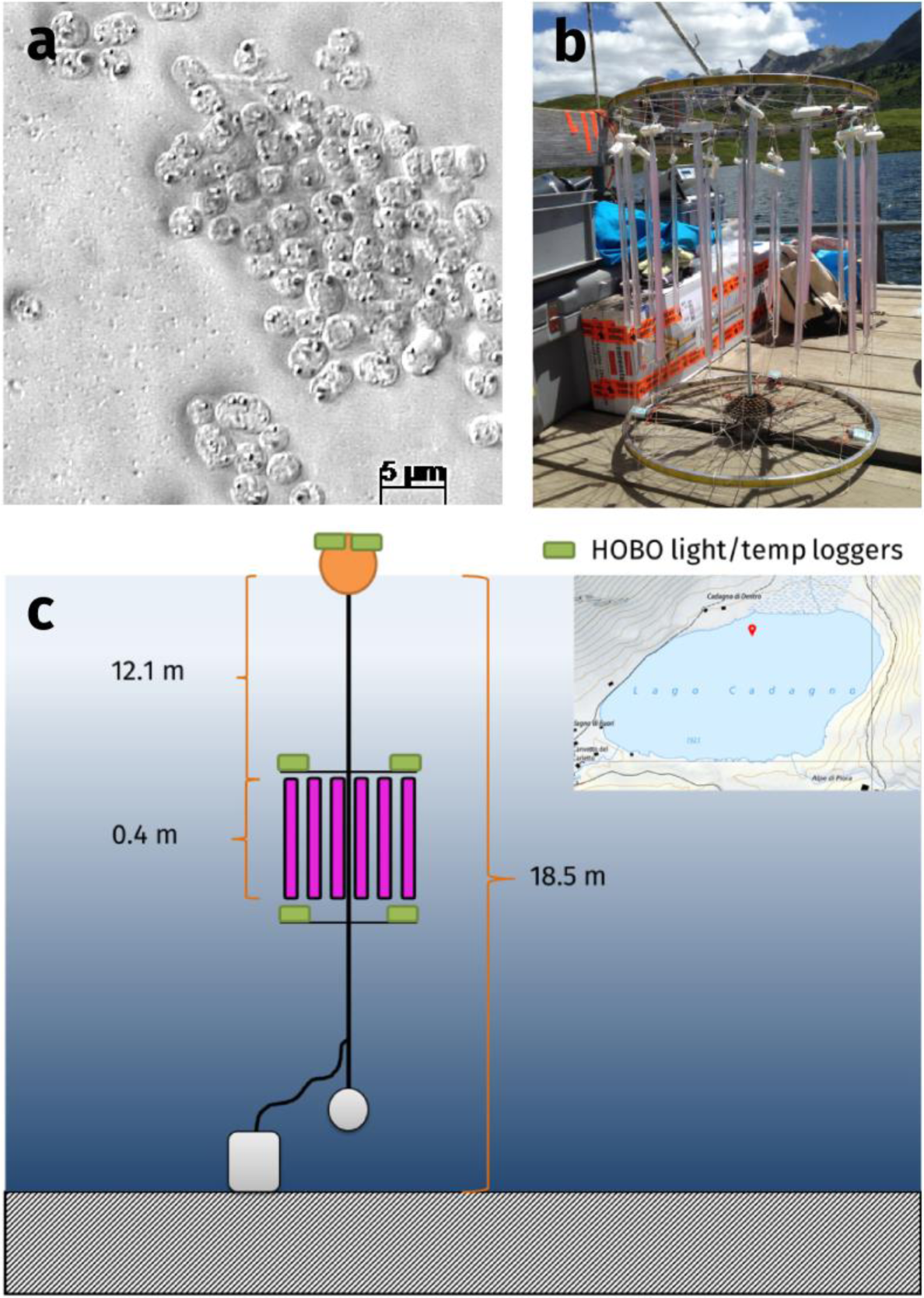
Depiction of strain “*Thiodictyon syntrophicum*” sp. nov. strain Cad16^T^ cells, sampling site and the experimental setup. **a)** Phase-contrast microscopic image of pure cultures of str. Cad16^T^ with sulfur inclusions visible as highly refractive particles. **b)** Set-up of the strain Cad16^T^ cultures in dialysis tubes attached to a support grid. **c)** Mooring scheme and location of the incubation experiment (initial depth, July to August) on Lake Cadagno. In pink indicated the dialysis tubes, in green the HOBO temperature and light sensors.

**Figure S2.**
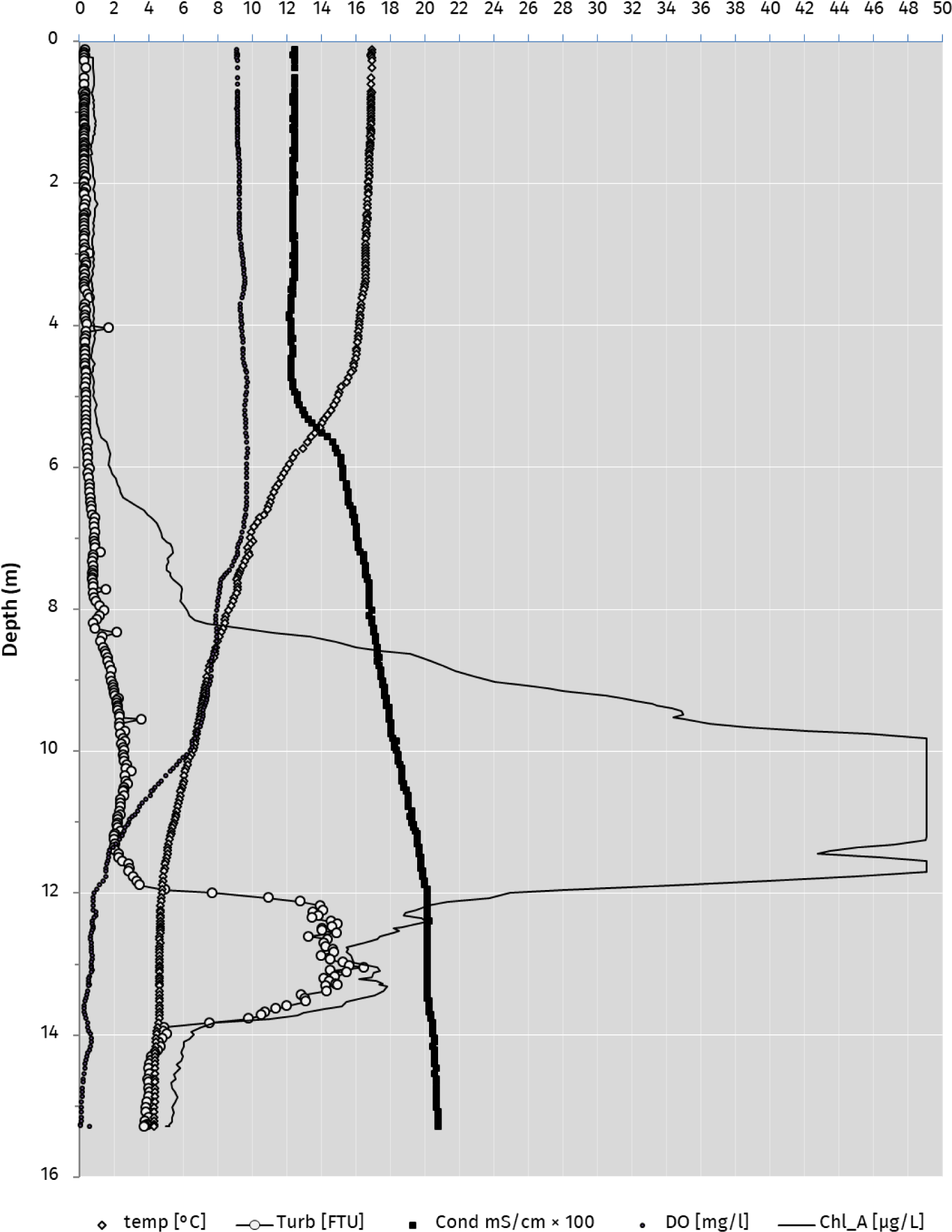
Conductivity, temperature, depth (CTD) profile on July 13 2017 at 2:44 pm above the deepest point of Lake Cadagno. Chlorophyll a (**––**), Temperature (◊), FTU: Formazin Turbidity Unit (○), Cond: Conductivity (▪), DO: Dissolved Oxygen (●),

**Figure S3.**
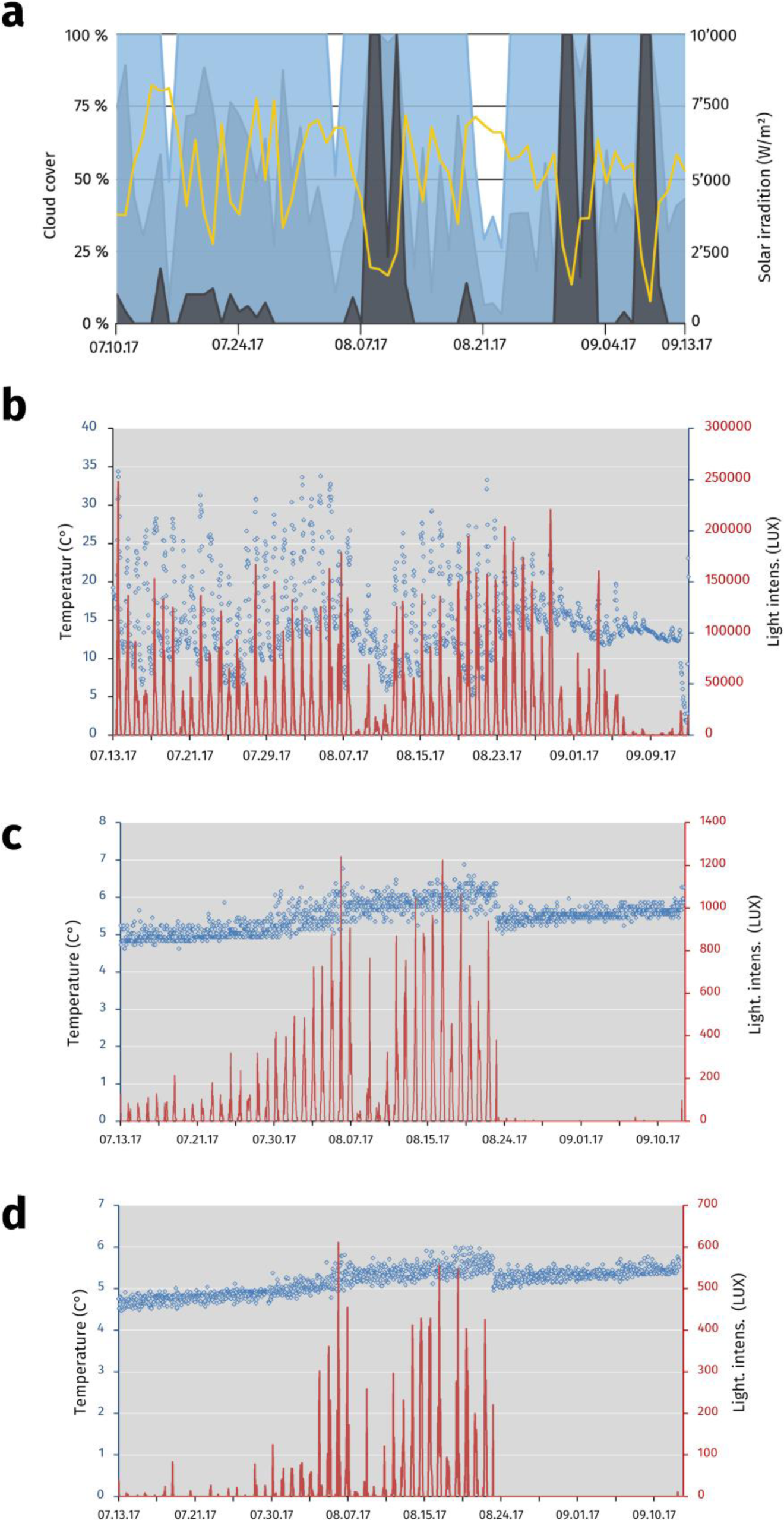
Meteorological data for the Piora valley and temperature and relative light availability at different depths of the mooring in Lake Cadagno from July 13 to September 13 2017. **a)** Sun light and cloud cover graph for the Piora valley from July 13 to September 13 2017. Data from meteoblue.com (yellow; sum of the daily shortwave radiation in W m^-2^, blue; maximal daily cloud cover in%, light blue; mean daily cloud cover in%, grey; minimal daily cloud cover in%) **b**) Temperature (black ovals) and average light available (red line) at the surface buoy of Lake Cadagno. The data logger was partly immersed in water, dampening the light and temperature readings (see values in August) **c)** A steady increase in average temperature and light availability from July to August is visible. The increase in the available light can be explained through the downward movement of photosynthetic bacteria over the season (see suppl. Fig. S5) Re-positioning of the rig on the August 23 2017 results in both, a drop in temperature, and light availability. Available light was reduced to an average of 0.4 µmol m^-2^ s^-1^. **d)** Low light availability and temperature are characteristic for the depth of around 12.4 m. As in c), temperature and available light values are steadily rising from July to August. No light was detected during daytime at 14.4 m depth after repositioning in August. Data was logged in hourly intervals for b)-d).

**Figure S4.**
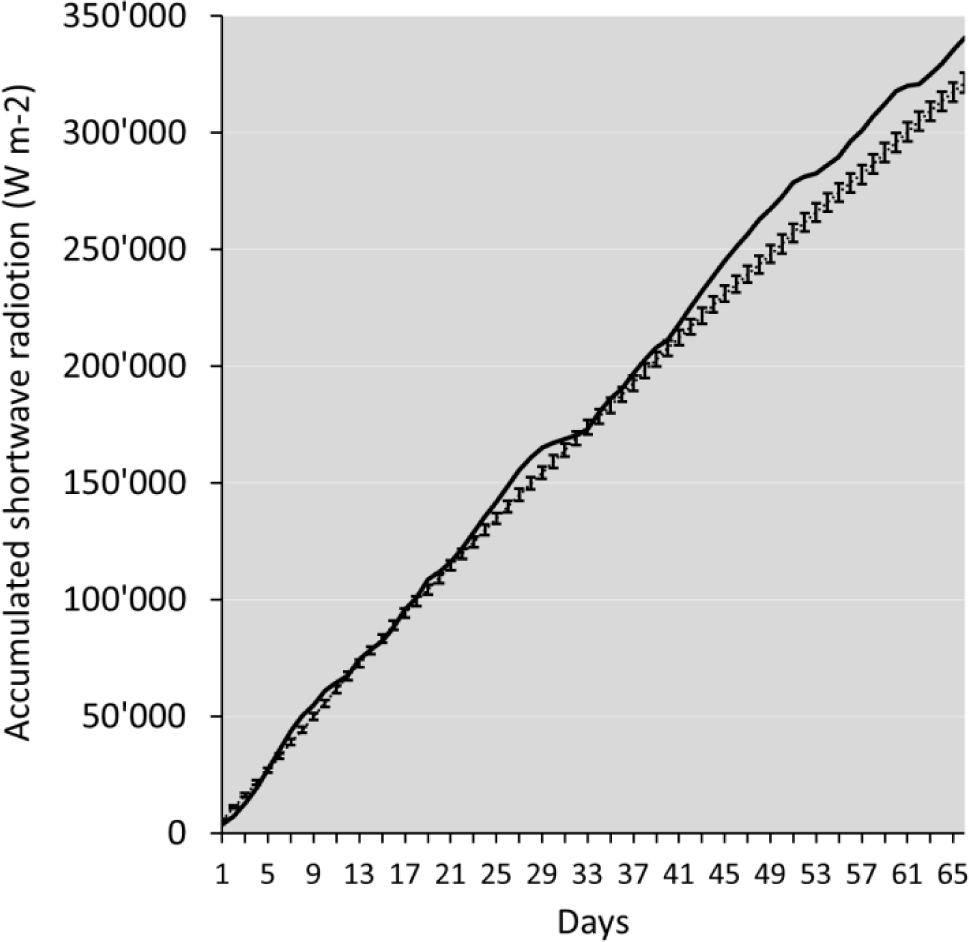
Accumulated surface shortwave radiation for 66 days from July 10 to September 12 2017 at Piora valley. The values for 2017 show above-average values (solid line). The mean values from 1985 to 2017 for the same time period are represented by the dotted line. Error bars indicate standard-error. Global radiation (diffuse and direct) on a horizontal plane given in Watt per square meter. Values derived from simulation data from meteoblue.com.

**Figure S5.**
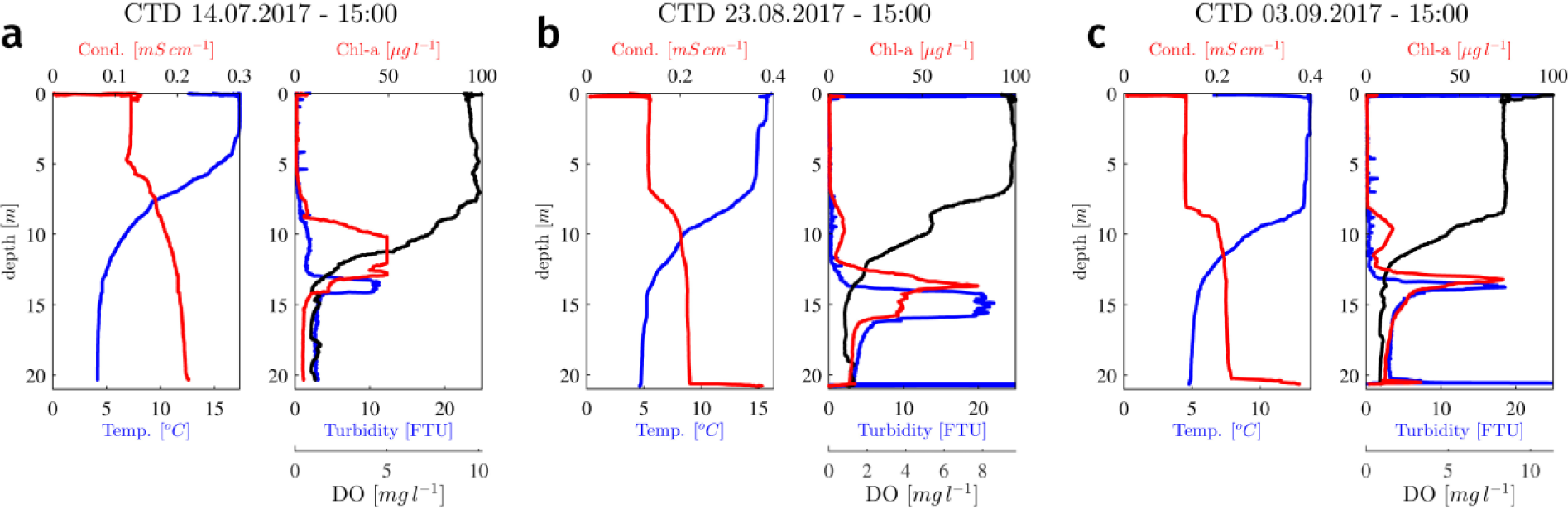
Vertical conductivity, temperature, depth (CTD) profiles from Lake Cadagno from July to September 2017 at the deepest point of the lake. Temperature, conductivity, dissolved oxygen, chlorophyll a and turbidity are displayed for: **a)** July 14 2017 **b)** August 23 2017 **c)** September 03 2017. Formazin Turbidity Unit (FTU). The data is courtesy of Dr. Oscar Sepúlveda Steiner, Dr. Damien Bouffard and Prof. Johny Wüest. Plots courtesy of Dr. O. Sepúlveda Steiner, APHYS Laboratory, EPFL, Switzerland.

**Figure S6.**
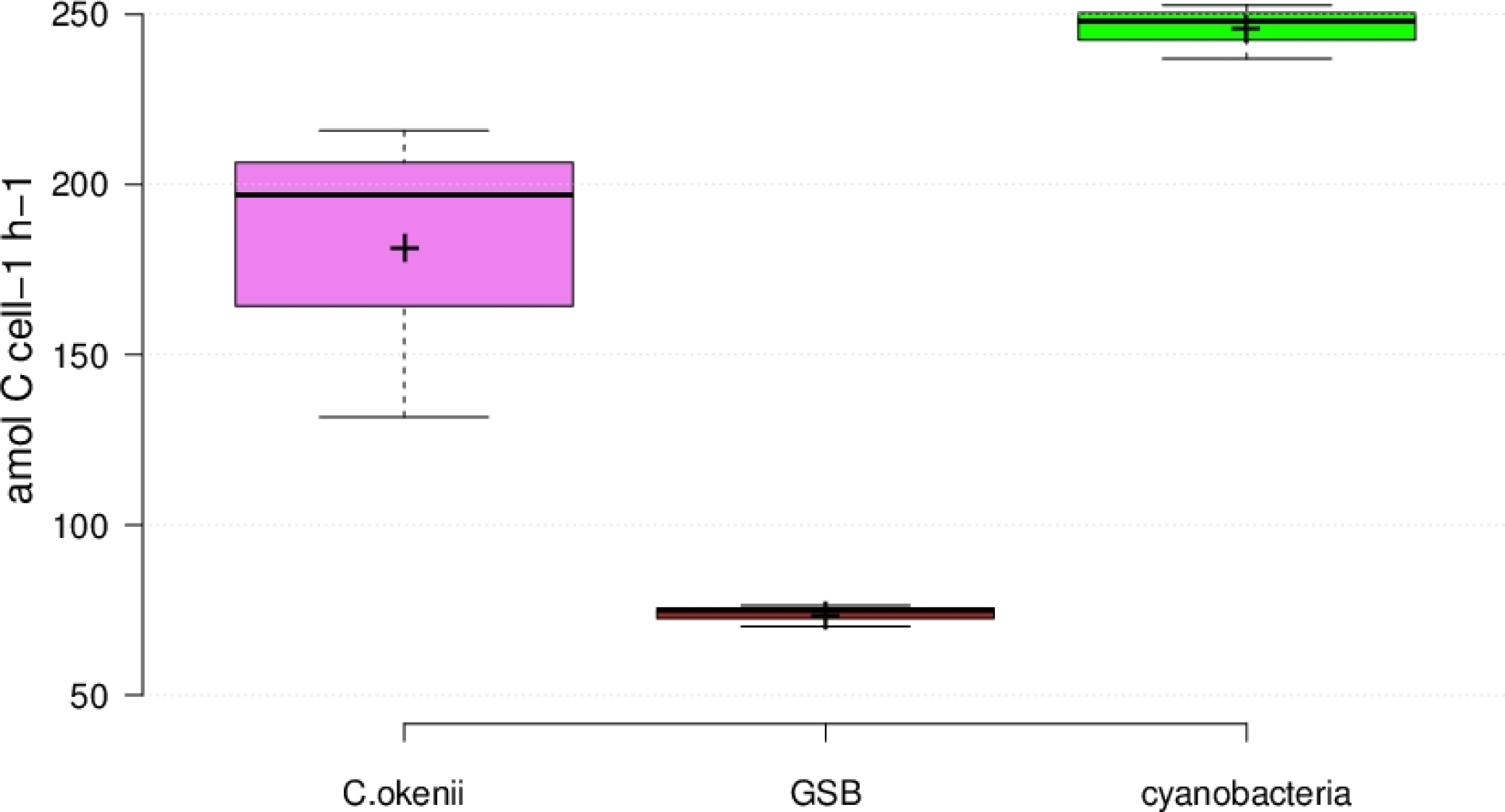
Average inorganic uptake rates at the chemocline of the three populations counted with FCM at 1:30 pm. *Chromatium okenii* (pink), green sulfur bacteria (GSB; brown) and cyanobacteria (green). Center lines show the medians; box limits indicate the 25^th^ and 75^th^ percentiles as determined by *R* software; whiskers extend 1.5 times the interquartile range from the 25^th^ and 75^th^ percentiles, outliers are represented by dots; crosses represent sample means. *n* = 3 sample points.

